# Temporal and spatial selectivity of hippocampal connectivity with sensorimotor cortex during individual finger movements

**DOI:** 10.1101/479436

**Authors:** Michael J. Miller, Douglas D. Burman

## Abstract

Cognitive control refers to brain processes involved in regulating behavior according to internal goals or plans. This study examines whether hippocampal connectivity with sensorimotor cortex during paced movements shows a pattern of spatial and temporal selectivity required for cognitive control. Functional magnetic resonance imaging activity was recorded from thirteen right-handed subjects during a paced, non-mnemonic (repetitive tapping) motor task. Direct and inverse connectivity in sensorimotor cortex were examined from psychophysiological interactions (PPI) from hippocampal seed activity during two sets of analyses: the first identified motor interactions relative to rest, whereas the second identified interactions in motor activity between fingers. Finger representations identified in a previous study were used to evaluate patterns of temporal and spatial selectivity in hippocampal connectivity. Changes in the magnitude of connectivity were identified within the sensorimotor representations of the first (index) through third (ring) fingers across time periods when each finger moved; at each finger representation, hippocampal connectivity was greatest when the represented finger was moving, reflecting temporal selectivity for the timing of finger movements. Similarly, the seeds associated with each finger representation differed in their magnitude of connectivity for adjacent finger representations, reflecting spatial selectivity for the moving finger. The patterns of spatial and temporal selectivity of connectivity during volitional movements in this study meets the criteria for cognitive control adapted from oculomotor studies.

## Introduction

To move purposefully, one or more signals within our brain direct the motor system to carry out the intended movement. This is the essence of cognitive control, the process by which goals or plans influence behavior. Although various regions of prefrontal cortex have been broadly implicated in cognitive control (1-5), the source of cognitive influence on the motor system is still subject to speculation.

Requirements for cognitive control of movements have been demonstrated in oculomotor studies, particularly those investigating the neural basis of reading (6-8) and other purposive eye movements (9-11). Such studies emphasize three components: 1) neural activity evident during conditions that require cognition; 2) spatial specificity, reflecting the spatial goal of the eye movement (i.e., target location); and 3) temporal specificity, which determines when the movement occurs.

The frontal eye field (FEF) is involved specifically in voluntary saccades, both in human and non-human primates (11, 12). Recording activity from nerve cells is generally avoided in humans, but FEF neurons in non-human primates reflect all three of these properties (13). These three properties are also evident from deficits following FEF lesions; for example, impairments in the accuracy and latency for a variety of volitional, but not reflexive eye movements are observed in both humans (11, 14-20) and non-human primates (21-24). In humans, lesions of the FEF and parietal eye field (PEF) differentially affect the selection of saccade targets and their timing (25); in monkeys, lesions of either area produce modest impairments in the accuracy and initiation of saccades, whereas combined lesions produce profound impairments (26). These studies demonstrate that the FEF, singly or in combination with PEF, is critically involved in the generation of volitional eye movements, providing the timing of purposive eye movements and their target location.

Analogous to the FEF in the oculomotor system (27), the primary motor cortex is necessary for volitional movements of individual fingers (28, 29), with short fiber tracts that connect postcentral with precentral regions of sensorimotor cortex (SMC) providing sensory feedback required for accurate performance (30). Neural activity in the FEF and sensorimotor cortex both reflect cognitive input. FEF properties that reflect cognitive input includes activity that predicts two sequential saccades (31-33), target selection (34-37), covert attention (38-42), and changes during learning (27, 43). Similarly, cognitive input to SMC is evident from neural changes during motor learning, both in its response properties (44-49) and changes in connectivity (50-52). Although cognitive input to both areas can be inferred, direct evidence for the source (or sources) of this input is sparse. Indirect evidence from neural properties and lesion effects has implicated the supplementary eye field (SEF) (53, 54), basal ganglia (55), and dorsolateral prefrontal cortex (56) in the cognitive control of eye movements, and the basal ganglia (57), cingulate (58), and prefrontal cortex (1, 59-63) for the cognitive control of skeletomuscular movements.

From this, it is tempting to speculate that the prefrontal cortex regulates finger movements. The prefrontal cortex has traditionally been associated with cognitive control, yet its connectivity to the SMC is indirect via the dorsal premotor cortex (2, 64, 65). Prefrontal areas are functionally coupled to premotor areas when learning movement sequences but not during repetitive tapping (66, 67), suggesting prefrontal cortex could be involved in cognitive control when learning complex movement sequences but not during simple movements. In its role, premotor cortex is analogous to the SEF (27), which is specialized for learning eye movement sequences (27, 68-70) and resolving conflicts between potential saccade targets (71, 72). Because prefrontal connectivity to FEF and SMC is indirect and limited to complex movements, cognitive control of simpler movements mediated specifically through FEF and SMC may arise from a different source. Thus, a relationship between neural activity and task complexity is unlikely when studying cognitive control in SMC, even though such a relationship has been used effectively to study cognitive control elsewhere (4, 73, 74).

The control of finger movements is more complex than eye movements for at least two reasons. First, the spatial reference for finger movements is different. Whereas eye movements are encoded from the current position of the eyes within the orbit (retinocentric space), finger movements are based upon the current position of the body (body-centered space); thus, the combination of finger muscles required to achieve a goal differs with different body positions. Second, the load on eye muscles is constant, whereas muscles moving the fingers may require different forces when moving objects of different resistance. When a task requires the initiation of finger movements from a set position with minimal load, however, requirements for cognitive control are the same as for eye movements: the neural mechanism must be active during volitional movements that require cognition, while specifying the timing and goal of the movement. The goal under these conditions reflects the body part that moves (e.g., hand or finger), requiring cortical specificity for neural activity within the SMC hand representation.

An earlier study examined hippocampal-SMC connectivity within the framework of volitional movements (75); the current study considers whether this connectivity shows spatial and temporal selectivity, necessary to specify which finger moves and when. Connectivity associated with movements of individual fingers is shown to be organized topographically, with connectivity in the representation of the moving finger differing from adjacent fingers (spatial selectivity); furthermore, connectivity within each finger representation is greatest when the finger moves, dropping when movement stops (temporal selectivity). Combined with the earlier study, hippocampal-SMC connectivity thus fulfills the three conditions of selectivity required for cognitive control. After considering alternative interpretations, we conclude that hippocampal connectivity is consistent with cognitive control of motor function, and describe experiments that could further elucidate this functionality.

## Materials and methods

### Subjects

Thirteen right-handed adults from the Chicago metropolitan area participated in the study (ages 24–59, mean=42.3, five females). Experimental procedures were explained to each subject before obtaining written consent; consent procedures complied with the Code of Ethics set forth in the Declaration of Helsinki, and were approved by the Institutional Review Board at the NorthShore University HealthSystem / Research Institute. Consented subjects had no history of the following: a previous concussion, psychiatric illness, learning disability, attention deficit disorder, abnormal physical or cognitive development that would affect educational performance, central neurological disease (or structural pathology), or neurosurgical intervention.

Immediately prior to their fMRI tests, subjects were trained on one cycle of each experimental task.

### Experimental tasks

The tasks were described in detail elsewhere (75). Briefly, the visual/motor task used in this study comprised 6 cycles of a specified sequence of visual and motor conditions. Motor conditions were preceded by an instruction screen, accompanied by a metronome ticking at 2 beats per second. During the sequence learning task, subjects rehearsed a remembered sequence of button presses, tapping in synchrony with the metronome; this was followed by a visual condition, which required fixation at the center without finger movements while a circular checkerboard flashed at 4Hz. This initial presentation of the visual task (“visA”) was followed by a block of repetitive tapping, subjects repeatedly tapped the same finger on both hands, corresponding to the button identified on-screen; the index finger (‘F1’) tapped first, and the finger increased incrementally every 4s (eight taps). After tapping movements from all four fingers (F1-F4 during time periods T1-T4, respectively), the visual condition was repeated (“visB”).

This report uses two sets of psychophysiological (PPI) connectivity for selectivity analyses; the first used the visA task condition as baseline (comparing activity during movement and rest), whereas the second used the T4 period (comparing activity between movement of different fingers).

### MRI data acquisition

Images were acquired using a 12-channel head coil in a 3 Tesla Siemens scanner (Verio). Visual stimuli projected onto a screen (Avotec Silent Vision) were viewed via a mirror attached to the head coil, and behavioral responses were recorded by Eprime (Psychology Software Tools, Inc.) from an optical response box (Current Designs, Philadelphia, PA). Blood-oxygen level dependent (BOLD) functional images were acquired using the echo planar imaging (EPI) method, using the following parameters: time of echo (TE) = 25ms, flip angle = 90°, matrix size = 64 x 64, field of view = 22cm, slice thickness = 3mm, number of slices = 32; time of repetition (TR) = 2000ms. The number of repetitions for the visual/motor condition was 182. A structural T1 weighted 3D image (TR = 1600ms, TE = 3.46ms, flip angle = 9 °, matrix size = 256 x 256, field of view = 22cm, slice thickness = 1mm, number of slices = 144) was acquired in the same orientation as the functional images.

### fMRI data processing

SPM12 software (http://www.fil.ion.ucl.ac.uk/spm) was used to process and analyze data, applying procedures described elsewhere (75,76). Briefly, images were spatially aligned to the first volume to correct for small movements, sinc interpolation minimized timing-errors between slices, and functional images were coregistered to the anatomical image and normalized to the standard T1 Montreal Neurological Institute (MNI) template, then smoothed with a 10mm isotropic Gaussian kernel. During analysis, conditions of interest were specified for sequence learning, visual, and each of the four fingers moving during the repetitive tapping block; global normalization scaled the mean intensity of each brain volume to a common value to correct for whole brain differences over time.

The representations for F1 (index finger) through F3 (ring finger) were adapted from an earlier study (77).

### Psychophysiological interactions (PPI)

#### Preprocessing

Connectivity analysis was carried out using psychophysiological interactions (75), modified to account for individual variability in connectivity (76). Each voxel in the left and right hippocampus of the normalized brain was sampled, as delimited by the aal atlas in the WFU PickAtlas toolbox (http://fmri.wfubmc.edu/software/PickAtlas).

To examine temporal and spatial selectivity, two sets of connectivity analyses were carried out, differing only in the interaction term used to identify task-specific activity. In the first analysis, an interaction term specified a greater effect of seed activity during movement of a single finger than during the visual condition; this identified the effect of movement relative to rest (e.g., T1>visA, T1 specifying the time period for tapping the first finger). In the second analysis, the interaction term specified a greater effect of seed activity during movement of one finger relative to another finger (e.g., T1>T4). Each set of analyses used a constant time period for the baseline comparison (visA and T4, respectively), so that differences in connectivity between finger representations or time periods would not result from different baselines. Following adjustments for regional differences in timing, a regression analysis identified the magnitude of the BOLD signal in SMC correlated with the PPI interaction term.

#### Seed selection

In this study, “functional seeds” were selected (see Burman 2019), i.e., the hippocampal voxel generating the greatest movement-related connectivity anywhere within SMC. Functional seeds were selected both for high values of direct connectivity within SMC (“MAX”) and for low values, which reflect inverse connectivity (“MIN”). For both direct and inverse connectivity, a voxel seed was selected from both the left and right hippocampus, and a conjunction (global) analysis was used to the magnitude of connectivity.

As noted elsewhere (75), the locus of maximal connectivity associated with functional seeds often lay outside the finger representations and could occur in either hemisphere. Seeds were not selected based on measurements within a specific finger representation.

#### Finger topography and spatio/temporal selectivity

A previous study used a variant of our paced repetitive tapping task to identify individual finger representations (77). This study used high-resolution functional images (1mm isovoxels), identifying a finger representation for the right hand as the left SMC region where activation during movement of one finger exceeded the mean activation during movement of the remaining fingers. The finger representation for the little finger (F4 as labeled in the current study) could not be mapped, perhaps because some subjects rotated their wrist rather than moving their little finger to press the key. The high-resolution activation map for each finger representation from the earlier study was superimposed on a low-resolution (4mm isovoxel) image used in the current study; low-resolution coordinates for each finger representation are recorded in Table 1. Activation maps were flipped along the x-axis to map finger representations in the right hemisphere. Because high-resolution activation did not coincide with the boundaries of the low- resolution voxels, a few voxels in the current study were associated with more than one finger representation.

**Table 1.** Finger representations in left and right sensorimotor cortex. Coordinates are shown based on 4mm isovoxels for the normalized brain; voxels that appear in more than one finger representation are shaded in colors to show the dual-finger representation.

| Left<br>postcentral | F1 | F2 | F3 |
| --- | --- | --- | --- |
|  | (-46,-24,54) | (-34,-32,58) | (-34,-32,54) |
|  | (-50,-24,54) | (-34,-32,62) | (-34,-32,58) |
|  | (-42,-24,58) | (-34,-36,62) | (-34,-36,58) |
| Left<br>precentral |  |  |  |
|  | (-38,-24,58) | (-30,-32,58) | (-34,-24,58) |
|  | (-42,-20,58) | (-30,-28,62) | (-34,-28,58) |
| Right<br>postcentral |  |  |  |
|  | (50,-28,54) | (34,-32,62) | (42,-32,54) |
|  | (42,-28,58) | (34,-36,62) | (38,-32,58) |
|  | (46,-28,58) | (34,-36,66) | (42,-32,58) |
|  | (38,-32,58) | (34,-32,58) | (38,-32,62) |
|  | (42,-32,58) | (38,-32,58) | (34,-32,58) |
| **Right**<br>**precentral** |  |  |  |
|  | (38,-20,54) | (34,-24,58) | (34,-24,54) |
|  | (42,-20,54) | (38,-24,58) | (34,-20,54) |
|  | (38,-20,58) | (34,-24,54) | (38,-24,54) |
|  | (38,-24,54) |  |  |

With the voxels identified for each finger representation (F1-F3), the amplitude of connectivity from each functional seed (S1-S3) was identified and plotted for each time period (T1-T4). Paired t-tests from each voxel within a finger representation across time periods provided a measure of temporal selectivity from a given seed; this identified significant differences in the magnitude of connectivity between two time periods, corresponding to the movement of different fingers. Similarly, paired t-tests between finger representations from the same seed provided a measure of spatial selectivity within a time period; this identified significant differences in the magnitude of connectivity among finger representations during each time period, when only one finger was moving. In this spatial analysis, the same number of voxels near the central representation of each finger were statistically compared, avoiding those voxels common to both fingers.

The absolute value of beta estimates for connectivity varied widely between subjects and analyses, reflecting subject variability in several factors tangential to the analysis of interest (e.g., subject variability in spatiotemporal offsets used to best fit SMC activity to the PPI interaction term, plus variability in response to the baseline condition). Results are reported as percentage changes in connectivity from the median amplitude across all conditions; the median amplitude was calculated for each voxel within a finger representation, with the total range specified separately for temporal and spatial selectivity analyses. A previous study showed connectivity changes selectively within the SMC hand representation (75); percentage changes in connectivity measures from the midline thus reflect the relative influence of seed activity on a finger representation during each condition.

## Results

### Temporal selectivity of connectivity within a finger representation

Temporal selectivity within a finger representation was demonstrated when the magnitude of connectivity was greatest during the time period that the represented finger was moving.

Evidence for temporal selectivity for functional seeds is shown in Figure 1 for both the L and R postcentral gyrus. In the L postcentral gyrus, the PPI interaction term compared seed activity during finger movements vs. the preceding visual rest condition; significant connectivity was not observed when comparing seed activity during movement of one finger relative to the fourth finger. In the R postcentral gyrus, the reverse was true; significant connectivity was observed only when the PPI interaction term compared seed activity during movement of one finger relative to activity during movement of the fourth finger.

**Figure 1.**
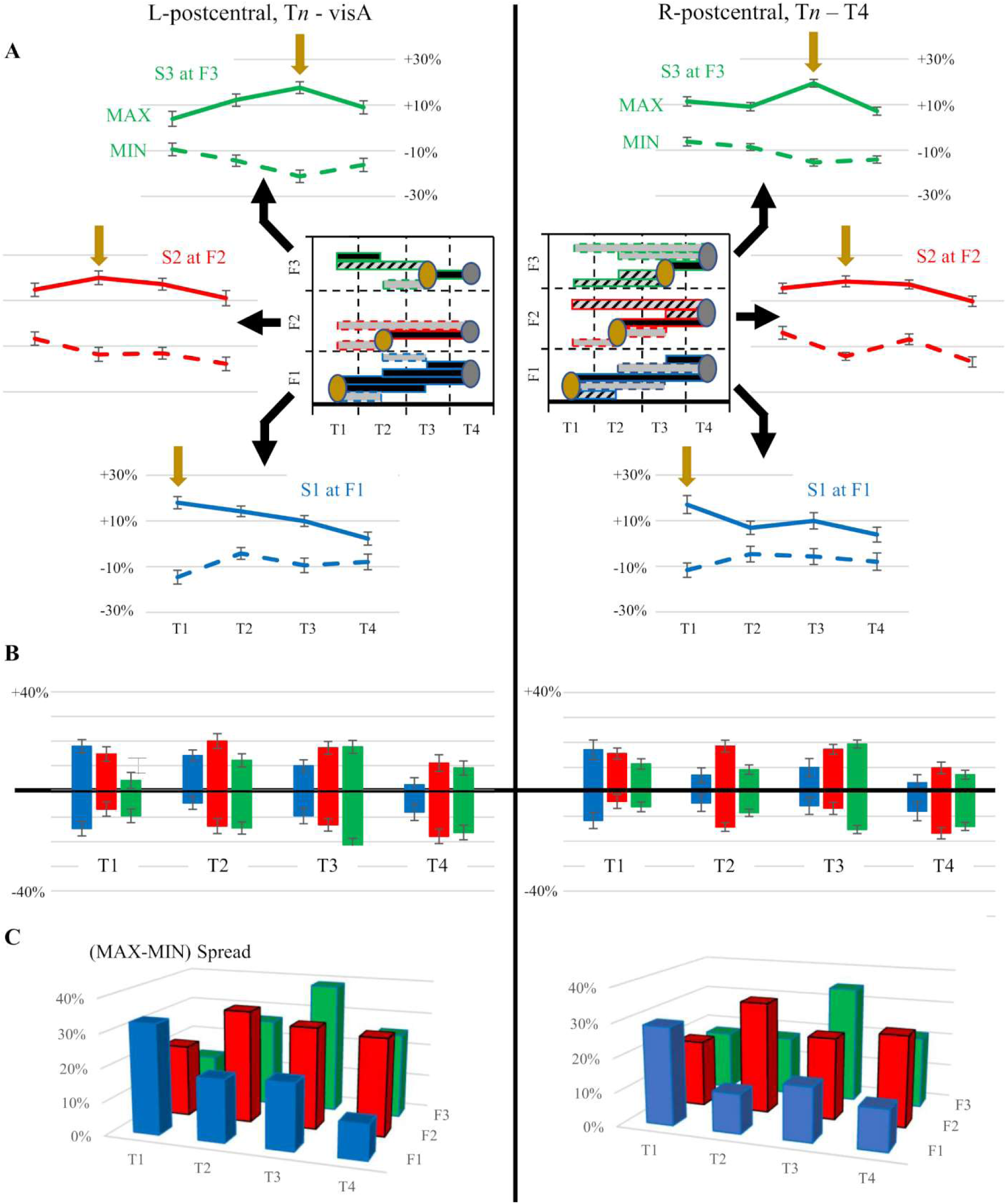
Temporal selectivity of hippocampal connectivity with SMC. A: For hippocampal functional seeds S1-S3, maximum and minimum connectivity with finger representation areas F1-F3, respectively, in the L and R postcentral gyrus are shown at time periods corresponding to the movement of fingers 1-4 (T1-T4, respectively). Gold arrows indicate the time period of movement for the fingers corresponding to F1-F3. Note that both maximum and minimum connectivity were greatest in each representation area during movement of its corresponding finger. Task-related connectivity in the postcentral gyrus was calculated relative to either the visual resting state VisA (L) or the movement of finger 4 (R). Error bars indicate the SEM. Insets: Paired t-test was used to identify significant differences in direct (gray) and inverse (black) connectivity among all time periods (T1-T4) at each finger representation F1-F3 in the L and R postcentral gyrus. Gold ovals indicate the period of movement for the fingers corresponding to areas F1-F3; silver ovals indicate the control condition for each side. In each finger representation, significant connectivity differences were typically linked to the period of movement for its represented finger and to the control state. B: Maximum and minimum hippocampal connectivity are compared for S1/F1 (blue) S2/F2 (red) and S3/F3 (green) at each time period T1-T4. During each period of movement, both the maximum and minimum connectivity were greatest in the moving finger’s representation. C: The Max-Min spread of connectivity is compared at each time period for S1/F1 (blue), S2/F2 (red), and S3/F3 (green). For each finger representation, the spread was greatest during the period of the represented finger’s movement.

Across all four time periods (T1-T4), Fig 1A shows the hippocampal connectivity maximum (MAX, solid line) and minimum (MIN, dashed line) for each functional seed (S1-S3) at its corresponding finger representation (F1-F3). In each hemisphere, MAX connectivity for a seed was greatest during the movement of its corresponding finger (e.g., S1 in F1 during T1); typically, the magnitude of MIN connectivity below the mean (inverse connectivity) was greatest during this same time period.

Hippocampal connectivity at a finger representation was compared between time periods using paired t-tests (Fig 1A, inset). In both hemispheres, connectivity during the time period where the represented finger was moving (demarcated by gold ovals) consistently differed from the control condition (gray ovals). Significant differences with the time period immediately before or after the movement of a given finger typically involved MIN connectivity (black bars extending one time segment from a gold oval). Significant differences for MAX connectivity (gray bars) was observed, however, at F3 in both hemispheres between T3 and T4, the control condition, and both MAX and MIN connectivity (striped bar) were observed at F1 in the right hemisphere between T1 and T2. MAX connectivity, however, more often produced changes spanning multiple time periods.

The MAX and MIN hippocampal connectivity values (Fig 1B) and the MAX-MIN spread (Fig 1C) were compared for each time period. The spread was greatest for each seed/representation area during the movement of its corresponding finger.

### Spatial selectivity across finger representations

In Figs 2-3, we examined the magnitude of connectivity within a specified time period for each seed (S1-S3), as well as differences in connectivity between finger representations. Fig 2 shows the connectivity analysis for the left postcentral gyrus, which used rest as the baseline; Fig 3 shows the same analysis for the right postcentral gyrus, which used the period of movement for F4 as the baseline (T4).

**Figure 2.**
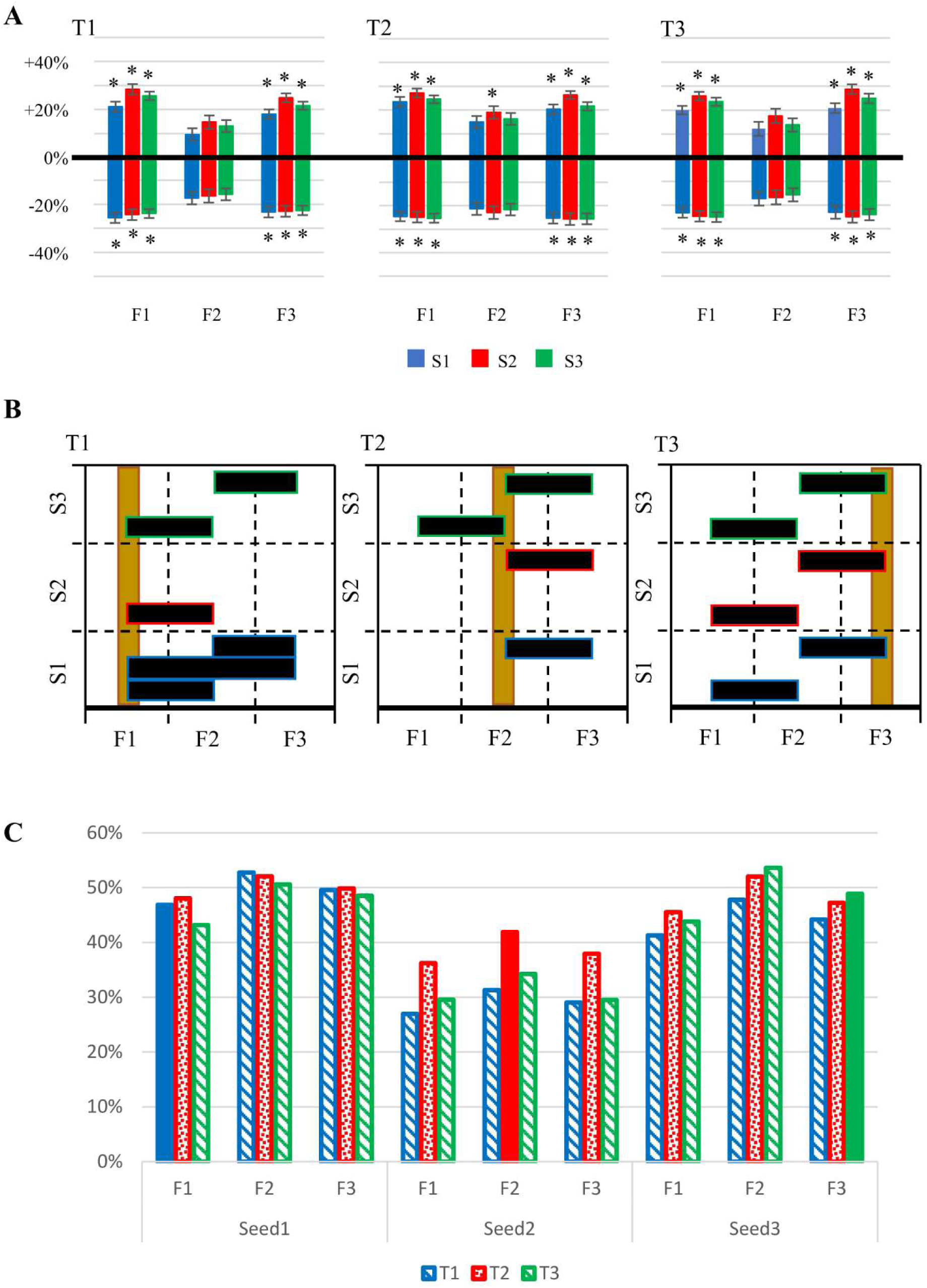
Spatial selectivity of hippocampal connectivity with L postcentral gyrus. Movement-related connectivity was calculated relative to the rest condition VisA. A: Magnitude of connectivity in finger representations F1-F3 for functional seeds during time periods T1-T3; both direct (MAX) and inverse connectivity values (MIN) are shown for each seed. MAX and MIN connectivity were both significant for F1 and F3, regardless of time period, whereas connectivity with F2 was significant only for the S2-MAX seed during T2. Error bars indicate the SEM, asterisks indicate significant connectivity after family-wise error correction (p<0.05). B: Paired t-test identified significant differences between finger representation during a time period; gold bars indicate the representation area corresponding to the finger moving during each time period. Significant differences were observed in the magnitude of inverse connectivity (black) between the representations of the moving finger and an adjacent finger. Differences in inverse connectivity were sometimes observed between other finger representations, but not direct connectivity (gray). C: Max-Min spread of connectivity was compared for each seed across time periods and finger representations; solid bars highlight the time period of movement for each finger representation. No consistent relationship was evident between the Max-Min spread and the timing of the finger movement associated with a finger representation.

**Figure 3.**
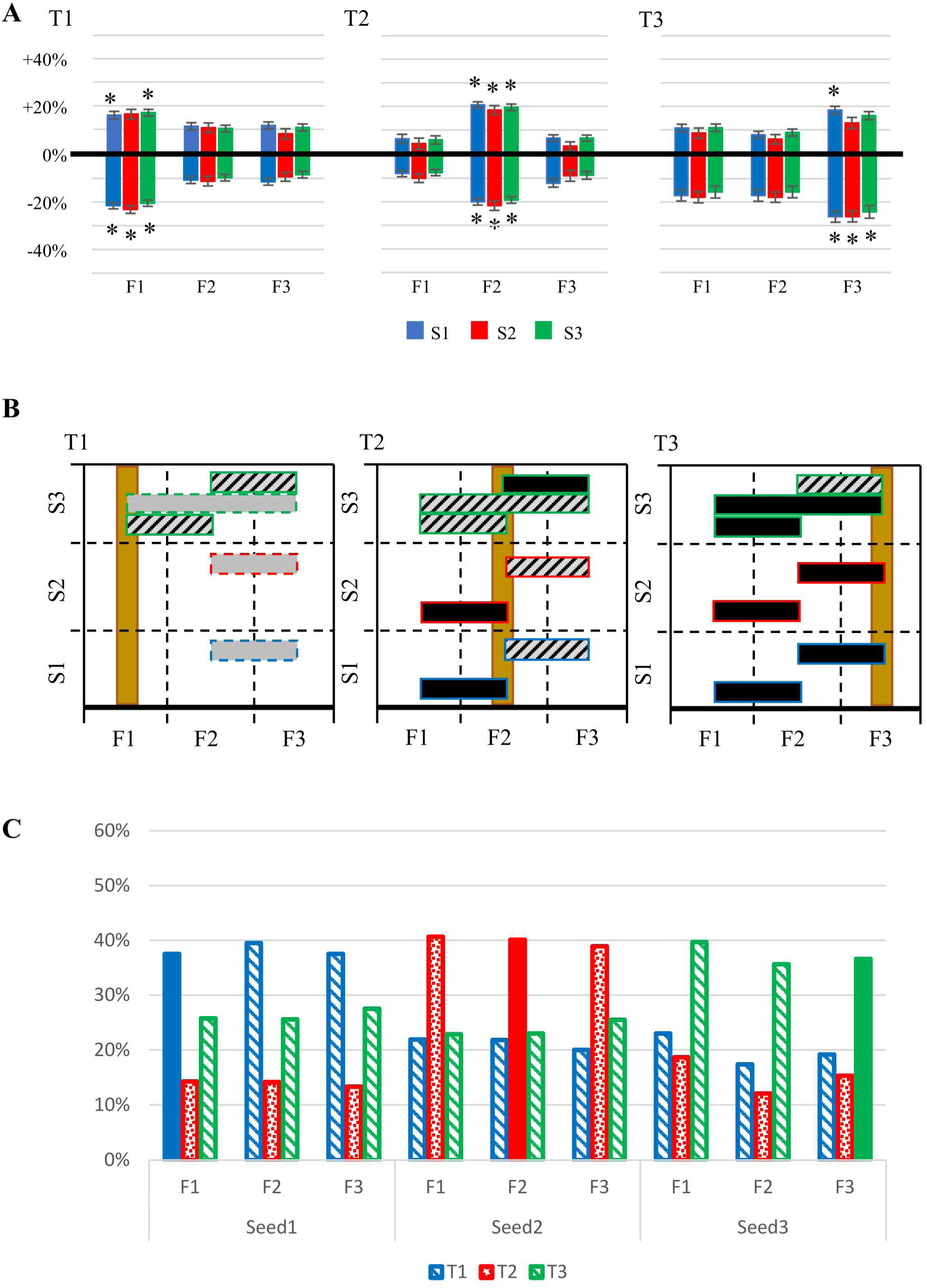
Spatial selectivity of hippocampal connectivity with R postcentral gyrus. Task-related connectivity was calculated relative to T4, the time period for moving finger 4; other conventions as in Fig 2. A: Maximum and minimum connectivity of hippocampal functional seeds. MAX and MIN connectivity were significant and larger within each finger’s representation when the represented finger was moving. B: Paired t-test identified significant differences in direct (gray) and inverse (black) connectivity between finger representations; striped bars indicate significant differences for both. Significant differences in connectivity were observed between the representations of the moving finger and an adjacent finger; during T1 and T3, significant differences were also observed between the non-moving fingers. Spatial differences in direct connectivity (silver) predominated during T1, whereas differences in inverse connectivity (black) predominated during T3. C: Max-Min spread of connectivity was compared for each seed across time periods and finger representations. All seeds provided the greatest MAX-MIN spread in connectivity during a single time period, regardless of the finger representation.

Fig 2A compares MAX and MIN connectivity within finger representations F1-F3 in the L postcentral gyrus during each time period (from T1 through T3). MAX and MIN seeds in F1 and F3 all showed significant connectivity in each time period, whereas significant connectivity in F2 was only observed for the S2 MAX seed during T2. The representation of the moving finger (i.e., F1 in T1, F2 in T2 and F3 in T3) did not necessarily show the greatest magnitude of MAX or MIN connectivity.

For each time period, Fig 2B shows differences in connectivity between finger representations; all significant differences involved inverse connectivity. The finger representation corresponding to the moving finger is demarcated by the gold bar. Hippocampal connectivity with the representation of the moving finger was significantly different from one or more neighboring fingers; connectivity differences between the stationary fingers were less consistent. Figure 2C compares the MAX-MIN spread at each finger representation across time periods. No consistent pattern in this spread was observed for any seed, either between time periods or between finger representations.

The pattern of MAX and MIN hippocampal connectivity for the R postcentral gyrus differed from the L postcentral gyrus (see Fig 3). Significant connectivity in each time period (Fig 3A) was present only in the finger representation for the moving finger. During T1 and T2, both MAX and MIN connectivity were significant when the represented finger was moving; during T3, only MIN connectivity (plus MAX for S1) was significant for F3.

During T1, only the S3 seed produced significant differences in F1 connectivity from that of other finger representations (Fig 3B). During T2 and T3, however, direct and/or inverse connectivity of the moving finger was significantly different from those of neighboring finger representations for all seeds. Also, the MAX-MIN connectivity spread was invariably highest in each finger representation when the represented finger was moving (Fig 3C).

### Topography and differentiation between fingers

In previous sections, T4 was used as the baseline for all connectivity analyses in the R sensorimotor cortex, which reflected differences in seed activity when different fingers are moving. Using visA as the baseline, by contrast, reflected differences in seed activity when a finger is moving vs. rest. This raises a question about the spatial resolution of hippocampal connectivity; can it differentiate between the movement of adjacent fingers?

Fig 4 shows the time course, extent, and topography of hippocampal connectivity delineated by differential seed activity from movement of adjacent fingers. The time course of the connectivity signal reflects the time course of the adjacent fingers compared; the time course when comparing activity from a finger’s movement with the preceding finger (top row) is earlier than for the same finger compared with the subsequent finger movement (bottom row). Although the amplitude of the connectivity signal may differ, the time course is similar from the left hippocampal seed (solid line) and the right hippocampal seed (dashed line). Connectivity associated with movement of each finger shows a topographical arrangement, with connectivity from F1 and F2 located inferior within postcentral gyrus to F4.

**Figure 4.**
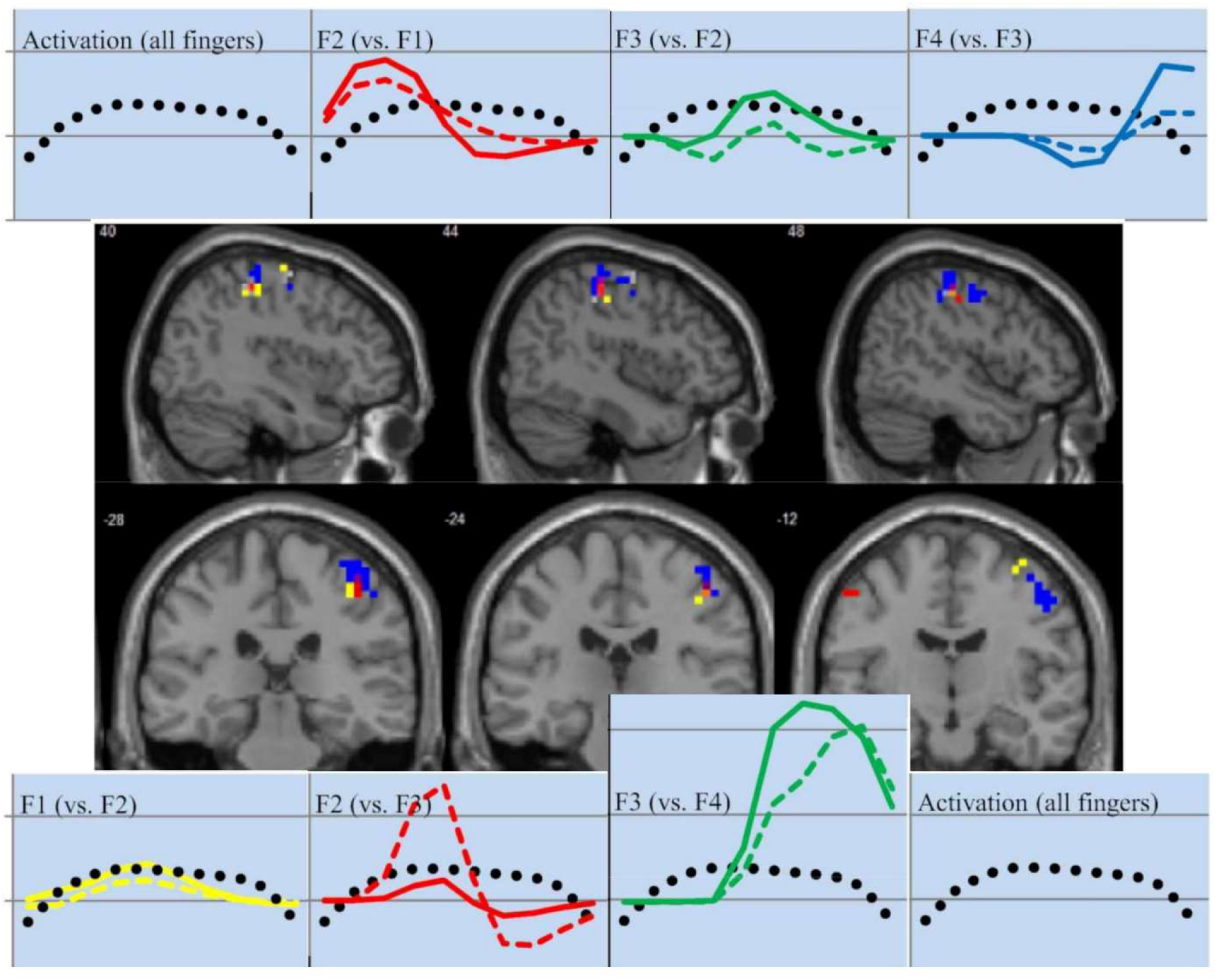
Hippocampal connectivity with R postcentral gyrus based on movements of adjacent fingers. Connectivity was calculated using functional seeds from the L (solid line) and R hippocampus (dashed line); functional activation across all time periods (dotted line) allows timing comparisons. Connectivity increases from the L and R hippocampal seeds followed the same time course, but connectivity derived from seed activity during movement of one finger vs. the preceding finger (top row) appeared earlier than connectivity derived from movement of the same finger compared to the successive finger (bottom row). Spatial overlap from the top and bottom analyses identified connectivity associated with a specific finger; connectivity associated with the fingers was organized topographically (middle row of images). F1=yellow, F2=red, and F4=blue.

The location, size, and magnitude of finger-specific connectivity is summarized in Table 2.

**Table 2.** Location of finger-specific connectivity in right SMC. Connectivity clusters in the right SMC from movement of individual fingers, surviving an intensity threshold of p=0.05 with a family-wise error correction

| **Finger/task interaction** | **Hemisphere (SMC)** | **Region** | **Cluster size** | **Z-score (peak)** | **Peak Coordinates** |
| --- | --- | --- | --- | --- | --- |
| F1>F2 | right | precentral | 9 | 4.36 | (38,-12,66) |
|  |  | postcentral | 18 | 4.05 | (38,-32,50) |
| F2>F1 | right | postcentral | 22 | 5.05 | (50,-24,50) |
|  |  | postcentral |  |  | (38,-32,62) |
|  |  | precentral |  |  | (50,-16,46) |
| F2>F3 | right | postcentral | 22 | 4.41 | (42,-28,54) |
|  |  | postcentral |  | 4.21 | (34,-32,58) |
| F3>F2 | right | postcentral | 58 | 5.11 | (50,-24,54) |
|  |  | precentral |  | 5.03 | (34,-12,66) |
|  |  | precentral |  | 4.69 | (46,-16,50) |
| F3>F4 | right | precentral | 58 | 4.01 | (34,-16,66) |
|  |  | postcentral |  | 3.61 | (50,-24,54) |
|  |  | precentral |  | 3.59 | (38,-16,46) |
| F4>F3 | right | precentral | 71 | 4.44 | (50,-12,46) |
|  |  | postcentral |  | 4.08 | (42,-28,58) |
|  |  | precentral |  | 4.07 | (38,-12,58) |

## Discussion

To evaluate the possible role of the hippocampus in cognitive control, this study sought to demonstrate spatial and temporal selectivity of hippocampal connectivity with the SMC during a sensorimotor task.

Previously we examined hippocampal-SMC connectivity during mnemonic vs. non-mnemonic finger movements and found connectivity increases during both tasks, suggesting a hippocampal involvement in volitional control (75). As noted in the introduction, cognitive control of a movement requires neural activity during volitional movements plus temporal and spatial selectivity.

### Temporal selectivity within the finger representations

Temporal selectivity for finger movements was identified as differences in the magnitude of connectivity across time periods within a given finger representation area. Consistent with this definition, maximal connectivity at each finger representation occurred when the finger was moving, with reduced connectivity during other time periods. A “spill over” effect was also evident, in which the time periods adjacent to the period corresponding to the moving finger exhibited elevated connectivity, likely because adjacent fingers do not move with complete independence.

Temporal selectivity was observed in both pre- and postcentral finger representations. This is expected, as motor and sensory systems operate jointly with movements closely tied to sensory feedback.

### Spatial selectivity

The topography of hippocampal connectivity with SMC was consistent with our previous study (77) and those of others (78-82), which showed overlapping activation between nearby fingers. Representations for later fingers (e.g., F4) were located dorsal to earlier fingers (F1-F3). The topographic organization of connectivity associated with individual finger movements followed this pattern in the postcentral gyrus of both hemispheres; although connectivity was present, no topography was apparent in the precentral gyrus.

During a single time period, spatial selectivity reflected differences in the magnitude of connectivity between finger representations. Within each time period, significant differences in spatial connectivity were observed; however, the L and R hemispheres exhibited different patterns of MAX and MIN connectivity, as will be summarized below. The functional relevance for many of these differences cannot be explained until we understand the specific behavioral effects mediated by MAX (positive) and MIN (inverse) connectivity.

In the L postcentral gyrus, the visual rest condition was required as a baseline to demonstrate significant effects. Significant MAX and MIN connectivity was observed for F1 and F2 during each time period, whereas F2 only showed greater MAX connectivity in its representation area during movement. The amplitude of MIN connectivity in the representation of the moving finger invariably differed from that of the adjacent finger, whereas no differences between fingers were observed for MAX connectivity. In the R hemisphere, by contrast, T4 was required as the baseline; furthermore, significant MAX and MIN connectivity changes were observed only in the representation of the moving finger. The magnitude of both MIN and MAX in the moving finger differed from those of an adjacent finger.

The complex spatial distribution of connectivity changes across finger representations in both hemispheres is not surprising, considering the multiple motor processes that occur during a single time period. Moving vs. waiting, activation of flexors vs. extensors, and movement readiness vs. actual movement are some of the opposing factors that can potentially result from direct vs. inverse connectivity. Fingers do not necessarily move independently of their neighboring fingers; moreover, deliberate movement of one finger may also involve the active suppression of stationary fingers.

In the L hemisphere, connectivity was measured relative to a baseline visual resting condition; the interaction term for this connectivity analysis provides a signal simply to “move this finger,” suggesting direct control of an individual finger independent of others. Connectivity in the R hemisphere, on the other hand, required an interaction term that explicitly signaled, “move this finger but not that one,” suggesting an interactive process between the control of different fingers. Because all subjects in this study were right-handed and volitional movements of the right hand are controlled by the left SMC, these findings suggest hippocampal influences on sensorimotor activity provide better independence of individual finger movements in the dominant hand.

Our analysis further demonstrated that differential connectivity was resolvable between adjacent fingers, although elevated connectivity for the moving finger could “spill over” into an adjacent finger representation. Significant differences in connectivity were observed during T1, for example, between the representations for F2 and F3, as well as between F1 and F2. A similar spillover effect has been observed for fMRI activation during finger movements, where movement-related activation extends into the representation of adjacent fingers (78-82).

### Hippocampal properties are consistent with cognitive control

Because the hippocampus is preferentially involved in conscious memories and environmental interactions (83, 84), studies of hippocampal interactions with the motor system have traditionally been focused on its role in explicit learning and memory consolidation. Hippocampal connectivity with the striatum increases during explicit motor learning and consolidation (85-87); consolidation after rehearsal is further reflected through increased motor activation following sleep in both the primary motor area and the hippocampus (85, 88). Our previous study showed that hippocampal connectivity to SMC occurs during paced repetitive tapping movements as well as sequence learning, and subjects’ anticipatory responses indicate volitional control of these movements (75). These findings demonstrate hippocampal involvement in movements and motor learning under explicit cognitive control; by contrast, the hippocampus is inactive during implicit learning of motor sequences (89). The second requirement for cognitive control is spatial selectivity. The hippocampus is sensitive to spatial location and navigation, with spatial functioning localized to the posterior hippocampus (90, 91), including space cells that encode a spatial map (92-96). Because the spatial location of the hand on the response pad was fixed in the current study, the spatial location of a button press corresponded to a SMC finger representation, and posterior structural seeds generated connectivity restricted to the SMC hand representation. Furthermore, spatial selectivity was observed between finger representations, as the magnitude of connectivity for the moving finger differed from that of the surrounding fingers from the representation of the finger currently moving. Thus, spatial properties of the hippocampus provided the selectivity in motor response required for cognitive control.

The third requirement for cognitive control is temporal selectivity. The hippocampus responds differentially to sequences of events that differ in order (97-100), and to the intervals between stimuli within a sequence (101-103). In the current study, cognitive awareness of the temporal intervals between metronome beats provided the basis for anticipatory behavioral responses. Furthermore, temporal selectivity was observed within each finger representation, such that connectivity was greatest when the represented finger was moving.

Thus, connectivity with SMC during motor tasks is consistent with known hippocampal properties and meets the three suggested criteria for the cognitive control of movements. This does not preclude an additional role in cognitive control of SMC from other areas. Prefrontal cortex, for example, has been suggested to play an indirect role through its connectivity with dorsal premotor cortex (2), although functional coupling of prefrontal with premotor areas is limited to movement sequences (66). During paced repetitive tapping, at least, hippocampal connectivity provides the best explanation for the cognitive control of SMC.

### Cognitive control vs. other theories of hippocampal function

There is currently no consensus about the functional significance of the spatial and temporal properties of the hippocampus. Some have suggested the hippocampus has cognitive functions beyond its traditional roles (104-106), whereas others suggest its diverse properties merely reflect the varied components of episodic and long-term declarative memory (92, 93, 107). Although hippocampal connectivity with SMC meets the requirements for cognitive control, could the results be better explained through its other known functions?

Memory of the pacing interval, plus associations between the numerical onscreen display and the corresponding fingers, were required for accurate performance during motor tasks in this study; these modest memory requirements might arguably require hippocampal input. Neither spatial nor temporal selectivity, however, is required to access this mnemonic information. Spatial selectivity is unnecessary because the mapping between numerals and fingers never changed, whereas temporal selectivity is unnecessary because widespread cortical rhythms (such as theta) could serve to time events. The observed pattern of spatial and temporal selectivity is required for cognitive control, however, to specify the finger to be moved and when.

An alternative explanation is that hippocampal connectivity in this study represents a form of sensorimotor processing, facilitating the transformation of sensory signals into motor output (108-111). This interpretation could explain why its topography and spatial selectivity were most prominent in the postcentral gyrus. Even without hippocampal intervention, however, extensive connections between pre- and postcentral gyrus provides sensory feedback (112), so aside from cognitive control, the role of hippocampal sensorimotor processing in SMC is unclear. Furthermore, facilitation of sensorimotor processing does not require temporal specificity. Given the directional influence of the hippocampus on SMC (implicit in PPI analysis), a hippocampal role in sensorimotor processing begs the question: what directs the hippocampus to provide relevant spatio-temporal information to SMC as needed for task performance? Considering the tight relationship between neural activity in the hippocampal seeds and SMC (shown in Burman 2019, Fig 2), an additional control signal in SMC seems unlikely.

This does not exclude the possibility of other sources of cognitive control under different conditions. By analogy with the oculomotor system, damage to a single structure (e.g., FEF or SMC) can produce deficits without eliminating all function (eye movements or hand movements) due to the preservation of other structures with overlapping functions (SEF / PEF, premotor cortex / basal ganglia). Similarly, executive functions show deficits following damage to prefrontal areas, yet are not eliminated (113-115). Hippocampal dysfunction in Alzheimer’s impairs volitional motor control (116), but with overlapping systems for cognitive control, it would not eliminate volitional movements.

### Implications and directions for future research

This study provides evidence that the hippocampus is involved in the cognitive control of volitional finger movements. Whether this represents a new function, or represents another component of a single cognitive function, is unclear. These alternative possibilities suggest two broad experimental questions for further investigation.

First, how extensive is the hippocampal role in cognitive control during volitional movements? Based on the criteria for cognitive control, hippocampal connectivity should be lost when “mindless” finger movements are generated while simultaneously performing a mentally- challenging task; connectivity also needs to be demonstrated during volitional movements of other body parts. In addition, a fixed spatial relationship between fingers and button presses in the current study allowed cortical specificity for finger movements to provide a measure of spatial selectivity; it’s unclear how the hippocampal pattern of cortical / spatial selectivity would be affected by changes in the spatial target of finger movements.

Second, is the hippocampal role in cognitive control limited to volitional movements? The hippocampus is critically involved in the formation and recall of episodic and declarative memories, thus fulfilling the first criterion for cognitive control (preferentially active during volitional states). Whether hippocampal interactions with cortical areas meet the remaining criteria for cognitive control over other mental functions is unclear, although theoretical accounts of memory function suggest they might. When spatial and temporal selectivity of hippocampal connectivity with cortical areas is necessary to perform cognitive tasks, a role of the hippocampus in cognitive control is implied. The role of the hippocampus in cognitive control may be evaluated through experimental manipulations that affect cognitive awareness and performance, evaluating effects of these variables on the connectivity of the hippocampus with brain areas directly involved in task performance. Comparing effects of these variables on hippocampal and prefrontal connectivity may help identify conditions where these regions are either complementary or synergistic in the cognitive control of behavior.

## Conclusions

The hippocampus showed task-specific connectivity with SMC during paced movement tasks, meeting the criteria for cortical control adapted from studies of cognition in the oculomotor system: associated with volitional movements, restricted to the representations of the moving fingers, and selective for the timing of the movements. Still present in a motor task without demonstrable learning or memory effects, this pattern of connectivity is consistent with known hippocampal properties, yet cannot be explained by its established functions. A role for the hippocampus in cognitive control is indicated; a collaborative role with prefrontal cortex is suggested from their close functional relationship across tasks and neurological conditions.

